# Bagremycin Antibiotics and Ferroverdin iron-chelators are synthetized by the Same Gene Cluster

**DOI:** 10.1101/631242

**Authors:** Loïc Martinet, Aymeric Naômé, Benoit Deflandre, Marta Maciejewska, Déborah Tellatin, Elodie Tenconi, Nicolas Smargiasso, Edwin de Pauw, Gilles P. van Wezel, Sébastien Rigali

## Abstract

Biosynthetic gene clusters (BGCs) are organized groups of genes involved in the production of specialized metabolites. Typically, one BGC is responsible for the production of one or several similar compounds with bioactivities that usually only vary in terms of strength and/or specificity. Here we show that the previously described ferroverdins and bagremycins, which are families of metabolites with different bioactivities, are produced from the same BGC, whereby the fate of the biosynthetic pathway depends on iron availability. Under conditions of iron depletion, the monomeric bagremycins are formed, which are amino-aromatic antibiotics resulting from the condensation of 3-amino-4-hydroxybenzoic acid with *p*-vinylphenol. Conversely, when iron is abundantly available, the biosynthetic pathway additionally produces a molecule based on *p*-vinylphenyl-3-nitroso-4-hydroxybenzoate, which complexes iron to form the trimeric ferroverdins that have anticholesterol activity. Thus our work challenges the concept that BGCs should produce a single family of molecules with one type of bioactivity, the occurrence of the different metabolites being triggered by the environmental conditions.

## Introduction

Specialized metabolites are natural products that play essential roles by helping the producing strain to cope with various stresses, used as weapons to outcompete neighboring commensals, or required at particular physiological or developmental stages [1, 2]. Besides their role in improving the fitness of microorganisms in their habitat, amongst others, molecules emanating from the so-called secondary metabolism are of foremost therapeutic and agro-industrial importance [3, 4]. These bioactive compounds are produced by a machinery encoded by a group of genes - a Biosynthetic Gene Cluster (BGC) – that, next to biosynthetic genes, typically includes genes for expression control, self-resistance and export [4–6].

Genome sequencing of secondary metabolite-producing microorganisms has revealed the enormous potential to increase the known chemical space [5], with the promise of new leads in human therapies or for sustainable agriculture. One of the drivers of the renewed interest in natural products (NPs) was the discovery of so-called cryptic BGCs that are silent under routine laboratory conditions and may therefore specify molecules that had so far been missed during pharmaceutical screening [7–9]. An approach that is rapidly gaining momentum is to express cryptic BGCs in a heterologous chassis strain or superhost, and change promoter elements within the BGC by those that are expected to result in high expression under laboratory conditions. However, there are several problems associated with this approach. Firstly, it is yet not very amenable towards high-throughput strategies, and it is hard to establish the promise of a certain BGC based on the DNA sequence alone [10]. Secondly, examples are accumulating of NPs that are produced from multiple BGCs, or by strains in coculture [11, 12]. Alternatively, a single BGC may also be responsible for the production of many (up to over 100) structurally related molecules that differ in terms of their activity [13–15], or BGCs can be associated as “superclusters” to produce one or more similar molecules [16, 17]. Thus, to optimally exploit the chemical space of NPs, we need to understand the connection between the genomic and chemical diversity of their biosynthetic pathways.

Here we provide an example that further illustrates the complexity to estimate the metabolome profile and the industrial potential of a microorganism based on the genomic information alone. Our work revealed that the ferrous compounds called ferroverdins (Figure 1, structures 1 to 3 [18–20]), which are inhibitors of cholesterylester transfer protein (CETP) and thus molecules potentially able to reduce the risk of atherosclerosis [21], and the antibiotics bagremycins (Figure 1, structures **4 to 9** [22–24]), are synthesized from the same BGC. These molecules not only have completely different biological functions, but their biosynthesis also requires different environmental triggers. These findings challenge the concept suggesting that one BGC is associated with the production of similar compounds in terms of building blocks, structure, bioactivity, and condition for production.

**Figure 1.**
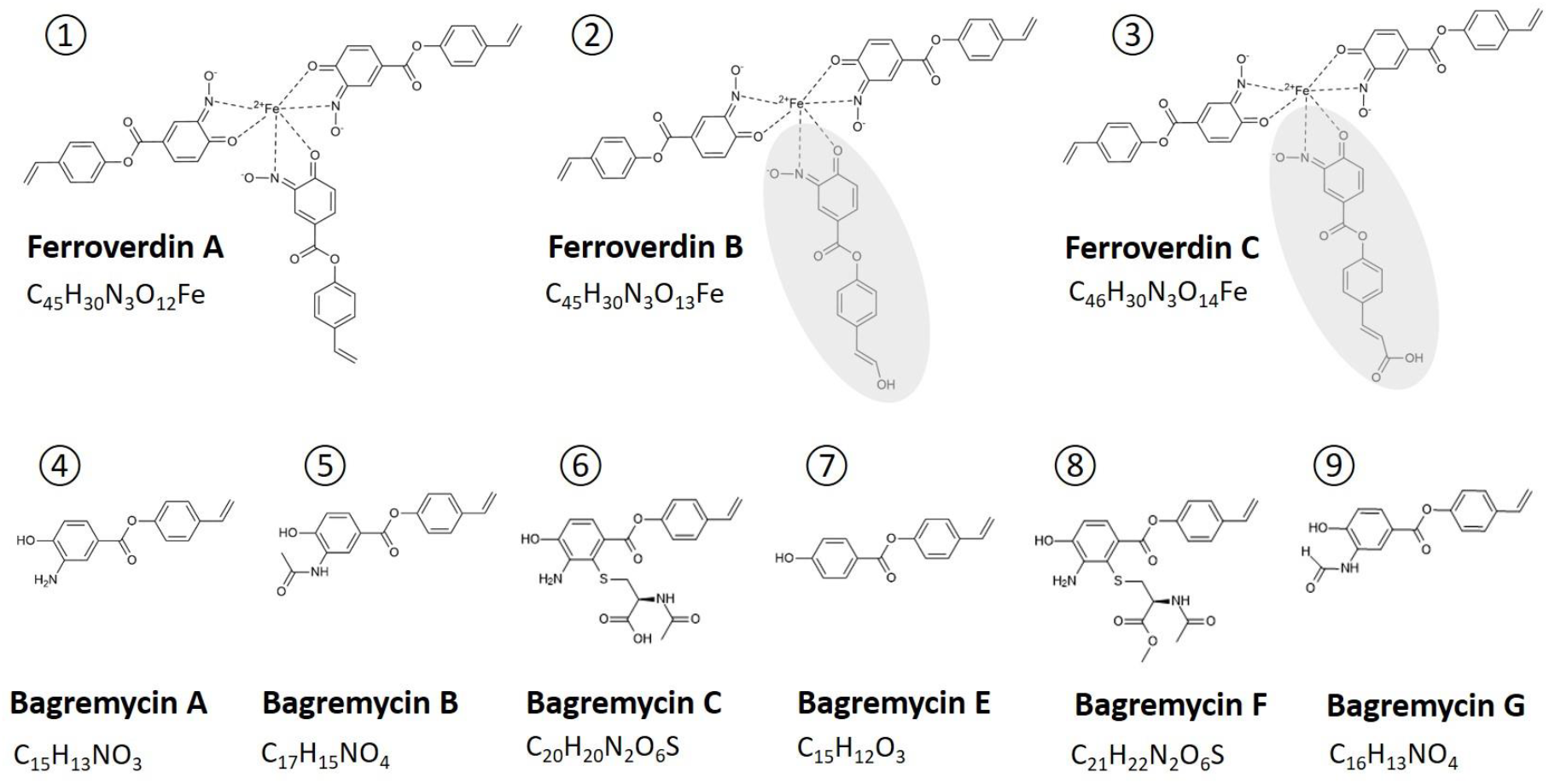
Structures of ferroverdins and bagremycins produced by *S. lunaelactis* strains. The top line displays the structures of ferroverdins (numbers 1 to 3) and the bottom line the structures of bagremycins (numbers 4 to 9), including their molecular formula. The monomer specific to ferroverdin B (hydroxy-*p*-vinylphenyl-3,4-NHBA), and the monomer specific to ferroverdin C (and carboxy-*p*-vinylphenyl-3,4-NHBA) are shaded in grey.

## Results

### *Streptomyces lunaelactis* produces ferroverdins and bagremycins

Strains of *Streptomyces lunaelactis*, including the type-strain MM109^T^, were previously isolated from cave moonmilk deposits [25–27]. The strain MM109^T^ produces a green pigment that was attributed to the biosynthesis of ferroverdin A [25], a homotrimer of *p*-vinylphenyl-3-nitroso-4-hydroxybenzoate (*p*-vinylphenyl-3,4-NHBA) complexed with one ferrous ion [18, 19]. To assess the metabolomic response of *S. lunaelactis* including conditions in which ferroverdin A is produced, strain MM109^T^ was grown under different culture conditions. For this, disc diffusion assays were performed, in the presence of a range of different metals.

This revealed that only the addition of FeCl_3_ triggered production of the green pigmented ferroverdin A by *S. lunaelactis* MM109^T^ (Figure 2A). Production of ferroverdin A (structure **1** in Figure 1, and Figure 2C) was triggered at a FeCl_3_ concentration as low as 0.01 mM (Figure 2B). UPLC-MS/MS analysis of extracts of strain MM109^T^ identified an ion peak of m/z 876.11 [M^-^], corresponding to ferroverdin B (structure **2** in Figure 1), and a distinct ion peak of m/z 904.11 [M^-^] corresponding to ferroverdin C (structure **3** in Figure 1). Ferroverdin B and ferroverdin C heterotrimers differ from ferroverdin A in only one of the three ligands used to complex ferrous iron, namely hydroxy-*p*-vinylphenyl-3,4-NHBA for ferroverdin B, and carboxy-*p*-vinylphenyl-3,4-NHBA for ferroverdin C (Figure 1).

**Figure 2.**
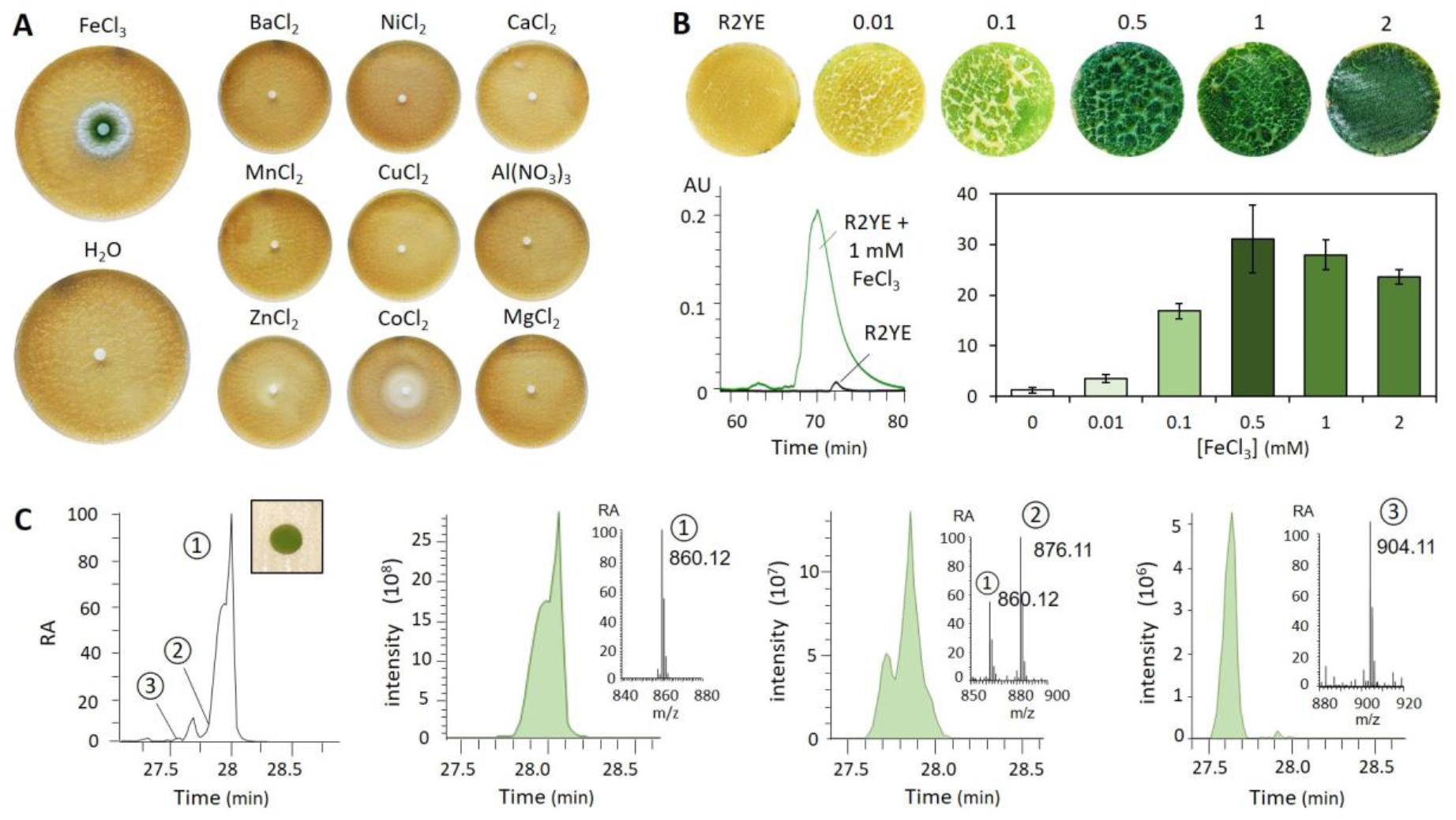
Production of ferroverdins by *S. lunaelactis* MM109^T^. **A.** Paper disc diffusion assays with various metal salts (1 mM). Note that FeCl_3_ is the only metal salt able to trigger the green pigmentation of the mycelium of *S. lunaelactis* MM109^T^ grown on R2YE agar plates. **B.** Induction of ferroverdin production by iron. Phenotype (top panels), HPLC profiles of the crude metabolite extracts (bottom left panel), and quantification (bottom right panel) of ferroverdin production by *S. lunaelactis* MM109^T^ grown on R2YE agar plates supplied with various concentration of FeCl_3_. The HPLC profiles performed with crude acetonitrile extracts of *S. lunaelactis* MM109^T^ grown on R2YE agar plates (black line) or R2YE medium supplied with 1mM FeCl_3_ (green line). **C.** Extracted ion chromatograms (EIC) of the three ferroverdins detected in the full extract of *S. lunaelactis* MM109^T^.

Under culture conditions allowing the production of ferroverdins, the crude extract of *S. lunaelactis* MM109^T^ displayed antibacterial activity against Gram-positive bacteria, as exemplified by the test strain *Staphylococcus aureus* (Figure 3A). As pure ferroverdins do not possess antibacterial activities against *S. aureus* (Figure 2C, left panel), we undertook the extraction, purification, and identification of the antibacterial metabolites produced by *S. lunaelactis*. HPLC fractionation and subsequent analysis of the active fractions by UPLC-MS/MS identified molecular ion species that corresponds to bagremycin A (m/z 254.0824 [M-H]^-^), bagremycin B (m/z 296.0931 [M-H]^-^), bagremycin C (m/z 415.0974 [M-H]^-^), bagremycin E (m/z 239.0714 [M-H]^-^), bagremycin F (m/z 429.1133 [M-H]^-^), and bagremycin G (m/z 282.0777 [M-H]^-^) (Figure 3BC, and structures **4, to 9** in Figure 1). Bagremycins are amino-aromatic antibiotics resulting from the condensation of 3-amino-4-hydroxybenzoic acid (3,4-AHBA) with *p*-vinylphenol [22–24].

**Figure 3.**
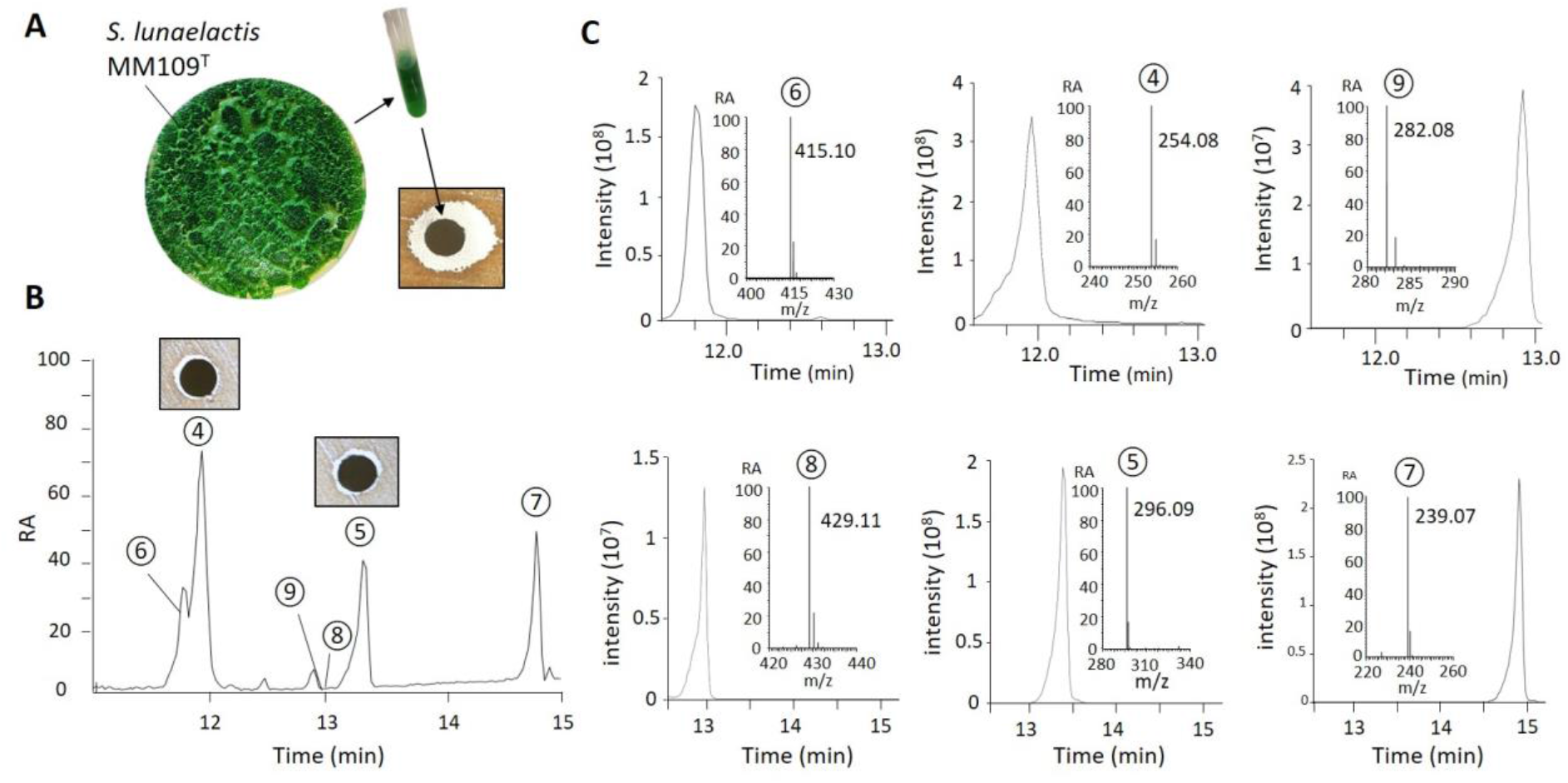
Production of bagremycins by *S. lunaelactis* MM109^T^. **A** Phenotype of *S. lunaelactis* MM109^T^ grown on R2YE + 1 mM FeCl_3_ agar plates, and antibacterial activity of its acetonitrile full extract (both intracellular and extracellular extracts). **B.** HPLC-separated fractions of the crude metabolites extract of *S. lunaelactis* MM109^T^. Note the antibacterial activity associated with pure bagremycin A (peak #4), and bagremycin B (peak #5). **C.** Extracted ion chromatograms (EIC) of the six known bagremycins detected in the full extract of *S. lunaelactis* MM109^T^.

MM109^T^ is the type strain of *S. lunaelactis*, but many other strains belonging to the species *S. lunaelactis* were isolated from different moonmilk deposits. MLSA analysis on 15 independently isolated *S. lunaelactis* strains branched them into different clusters (Figure 4A), and one representative of each cluster was investigated for the production of bagremycins and ferroverdins. The crude extracts of the selected eight different representative strains (MM25, MM31, MM37, MM40, MM83, MM91, MM109, and MM113) were analyzed by HPLC, which revealed that all strains except strain MM91 were able to produce at least one bagremycin and one ferroverdin (Figure 4B). These data suggest that the production of bagremycins, like the production of ferroverdins, is a common feature of moonmilk-dwelling strains belonging to the species *S. lunaelactis*.

**Figure 4.**
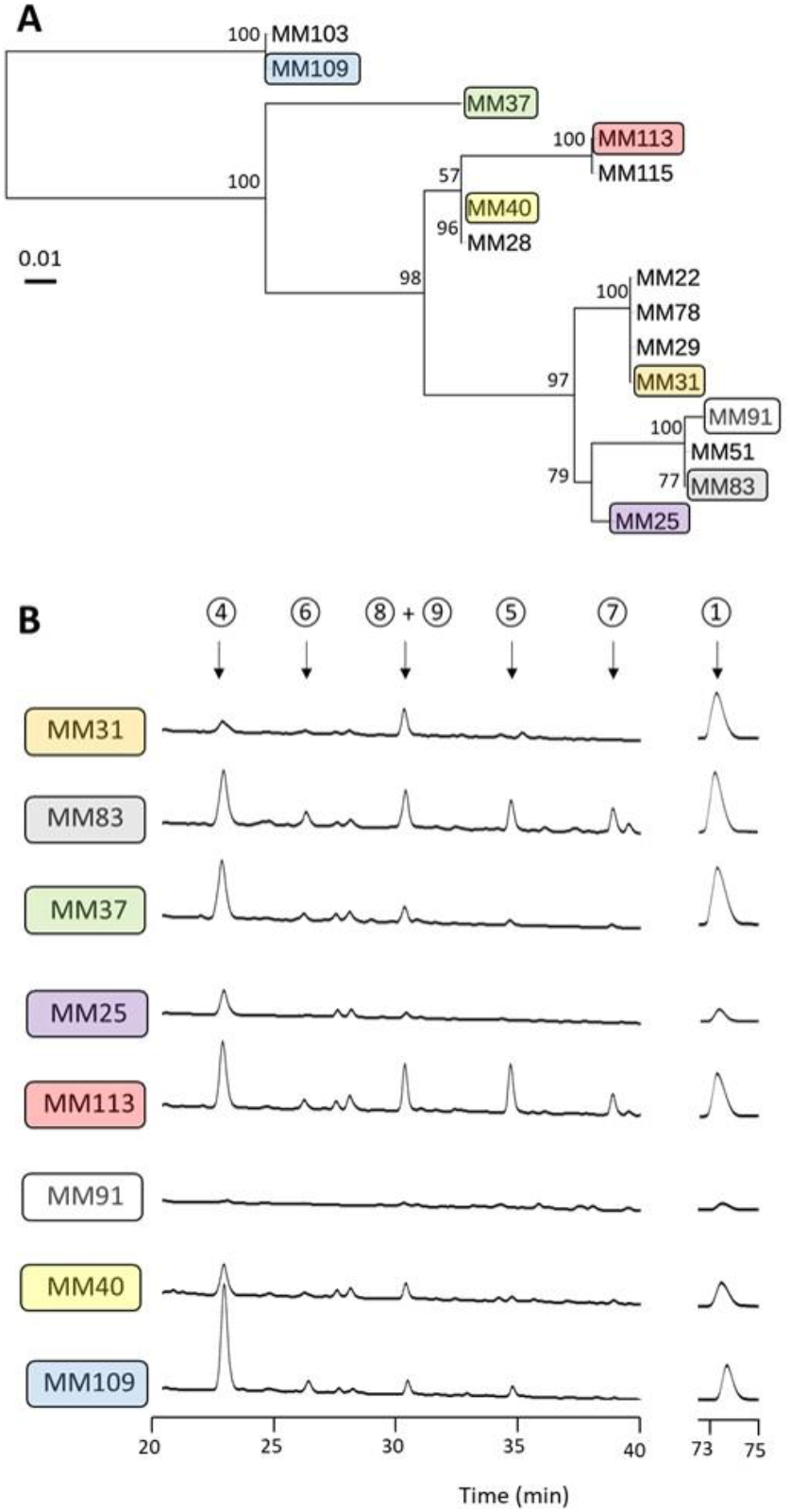
Production of ferroverdins and bagremycins by different *S. lunaelactis* strains. **A.** Multi locus sequence analysis (MLSA) of moonmilk isolates belonging to the species *S. lunaelactis*. **B.** HPLC profiles at retention times for monitoring bagremycin and ferroverdin production by eight different *S. lunaelactis* strains grown on the ISP7 medium.

### Identification of a BGC similar to *fev* and *bag* clusters in *S. lunaelactis* MM109^T^

In order to identify the genes involved in the production of bagremycins (*bag*) and/or ferroverdins (*fev*) in *S. lunaelactis* MM109^T^, the genome of this strain [28] was mined using the antiSMASH v4.0 software [29]. 37 BGCs were identified, namely 36 BGCs on the linear chromosome and one additional BGC on the linear plasmid pSLUN1 (Supplementary Figure S1). BGC #12 (from SLUN_21350 to SLUN_21430 [28]) shares strong gene synteny with the bagremycin BGC from *Streptomyces* sp. Tü 4128 [30, 31], with high similarity between the predicted gene products (coverage 99%, average identity 86.2%, average similarity 91.9%) (Figure 5A, Table 1). Surprisingly, BGC #12 shares similarly strong gene synteny and high similarity between the gene products with the *fev* cluster (GenBank: AB689797) of the ferroverdin producer *Streptomyces* sp. WK-5344 (coverage 99%, average identity 85.8%, average similarity 91.3%) (Figure 5A, Table 1). The level of identity between the proposed *fev* cluster of *Streptomyces* sp. WK-5344 and the *bag* cluster of *Streptomyces* sp. Tü 4128 are even greater with on average 96% aa identity and 97.4% similarity (Figure 5A, Table 1). The exceptionally high levels of average amino acid identity suggest that the biosynthesis of bagremycins and ferroverdins might be mediated by the same BGC. This was further supported by comparative analysis of the BGCs involved in the production of other amino/nitroso-aromatic metabolites *i.e*., the *nsp* gene cluster for biosynthesis of 4-hydroxy-3-nitrosobenzamide in *Streptomyces murayamaensis* [32], and the *gri* gene cluster involved in grixazone production is *Streptomyces griseus* [33]. The *nsp* and *gri* clusters share seven homologous genes with those identified in the *bag/fev* cluster (Figure 5A, Table 1), namely those encoding (i) the SARP-family transcriptional activators (BagI, FevR, NspR, and GriR), (ii) the LuxR-family transcription regulators (BagY, FevT, NspT, and GriT), (iii) the 3,4-AHBA synthases (BagC, FevH, NspH, and GriH), (iv) the DhnA-type aldolase (BagB, FevI, NspI, and GriI), (v) the o-aminophenol oxidases (BagH, FevF, NspF, and GriF), (vi) the copper chaperones (BagZ, FevE, NspE, and GriE), and (vii) the FAD-dependent oxygenases (BagK/G. FavA1/A2, NspA, and GriA). Phylogeny analyses revealed that Fev and Bag proteins always branch together in clusters separated from Nsp and Gri proteins (Figure 5B).

**Figure 5.**
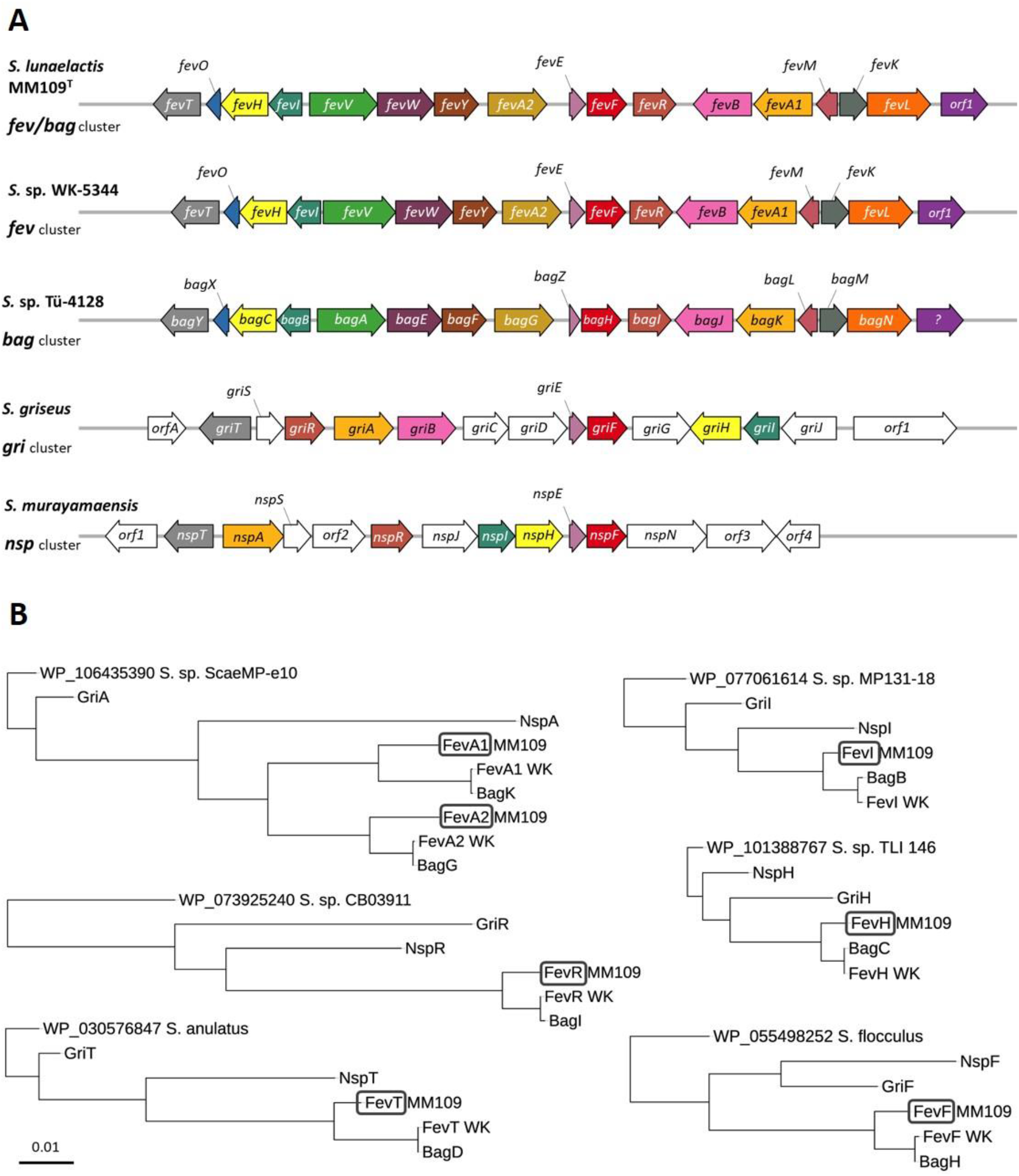
Comparative analysis of known BGCs involved in the production of nitroso- and amino-aromatic metabolites in *Streptomyces* spp. **A.** Genetic organization of the BGCs involved in ferroverdin (*fev*), bagremycin (*bag*), 4-hydroxy-3-nitrosobenzamide (*nsp*), and grixazone (*gri*) production is *Streptomyces* spp. See Table 1 for the known and/or predicted function associated with the product of each gene. **B.** Phylogeny analysis of proteins conserved among the ferroverdin (*fev*), bagremycin (*bag*), 4-hydroxy-3-nitrosobenzamide (nsp), and grixazone (*gri*) BGCs. The trees were rooted by including as outgroup the proteins most similar to *S. lunaelactis* MM109^T^ proteins.

**Table 1.**
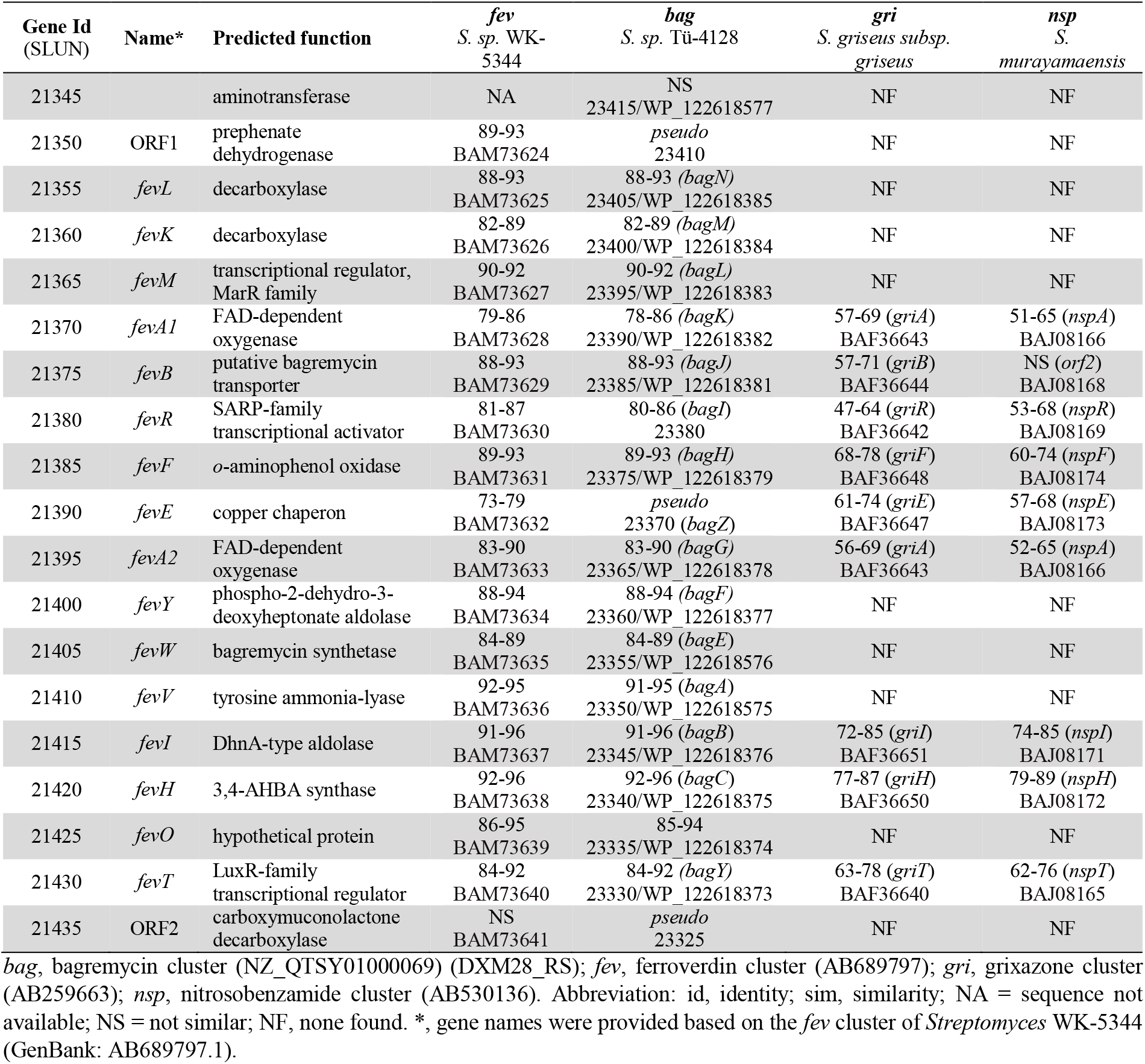
Elements of the ferroverdin/bagremycin BGC in *S. lunaelactis* and comparative analysis with similar clusters in other *Streptomyces* species.

**Table 2.**

| Plasmids and strains | Description | Source or reference |
| --- | --- | --- |
| **Plasmids** |  |  |
| pJET1.2/blunt | *E. coli* plasmid used for high-efficiency blunt-end cloning of PCR products (Amp<sup>R</sup>) | Thermo Scientific |
| pBDF019 | pJET1.2 derivative containing the *slun21380* (*fevR*) and its upstream (217 bp) and downstream (360 bp) regions (Amp<sup>R</sup>) | This study |
| pBDF020 | pJET1.2 derivative containing the *slun21380* (*fevR*) and its downstream (360 bp) region (Amp<sup>R</sup>) | This study |
| pSET152 | Integrative vector transmissible by conjugation from *E. coli* to *Streptomyces* spp. (*lacZa*, *ori* (pUC18), *aac(3)IV* (Apra<sup>R</sup>), *oriT* (RK2), *attP* (ØC31), *int* (ØC31)) | [42] |
| pBDF021 | pSET152 derivative containing the insert of pBDF019 cloned in *EcoRI* and *XbaI* sites (Apra<sup>R</sup>) | This study |
| pBDF027 | pJET1.2 derivative containing an internal fragment (791 bp amplified with x and x primers) of the *slun21380* (*fevR*) coding sequence (Amp<sup>R</sup>) | This study |
| pSET151 | Non-replicating plasmid in *Streptomyces* spp. (*lacZa*, *oriT* (RK2), *xylE*, *tsr* (Thio<sup>R</sup>), *bla* (Amp<sup>R</sup>), *ori* (pUC19)) | [42, 43] |
| pBDF028 | pSET151 derivative containing the insert of pBDF027 cloned in *XbaI* and *PstI* sites (Amp<sup>R</sup>, Thio<sup>R</sup>) | This study |
| pUZ8002 | Non-transmissible plasmid supplying transfer functions for mobilization of *oriT*-containing vectors from *E. coli* to *Streptomyces* spp. (Kan<sup>R</sup>) | [41] |
| pAU3-45 | pSET152 derivative with a thiostrepton resistance gene inserted into the blunted *NheI* restriction site (*lacZa*, *ori* (pUC18), *aac(3)IV* (Apra<sup>R</sup>), *oriT* (RK2), *attP* (ØC31), *int* (ØC31), *tsr* (Thio<sup>R</sup>)) | [44] |
| ***E. coli* strains** |  |  |
| DH5α | General cloning host | Gibco-BRL |
| ET12567 | Non-methylating (*dam*<sup>-</sup>, *dcm*<sup>-</sup>, *hsdS*<sup>-</sup>) host for transfer of plasmid DNA into *Streptomyces* spp., used together with pUZ8002 (Cml<sup>R</sup>, Tet<sup>R</sup>) | [45] |
| ***Streptomyces lunaelactis* strains** |  |  |
| MM109<sup>T</sup> | Wild-type and reference strain of *Streptomyces lunaelactis* | [25] |
| MM91 | Wild-type strain *Streptomyces lunaelactis* MM91 | [27] |
| MM109BD1 (*fevR*<sup>+</sup>) | MM109<sup>T</sup> derivative with pBDF021 inserted to add an additional copy of *slun21380* (*fevR*) controlled by its native promoter (Apra<sup>R</sup>) | This study |
| MM109BD2 (*fevR*<sup>-</sup>) | MM109 derivative with pBDF028 inserted to disrupt the gene *slun21380* (*fevR*) (Thio<sup>R</sup>) | This study |
| MM109BD3 | Complementation of strain MM109BD2 (*fevR*<sup>-</sup>) (Thio<sup>R</sup>, Apra<sup>R</sup>) | This study |
| SLUN pAU3-45 | MM109 derivative with pAU3-45 inserted as integration and thiostrepton resistance control (Thio<sup>R</sup>, Apra<sup>R</sup>) | This study |

Amp<sup>R</sup>: Ampicillin resistance; *lacZa*: LacZ galactosidase alpha subunit coding sequence; *ori* (pUC18/pUC19): Origin of replication of the plasmid pUC18/pUC19 (*E. coli*); *aac(3)IV*: Apramycin resistance marker; Apra<sup>R</sup>: Apramycin resistance; *oriT* (RK2): Origin of conjugative transfer from plasmid RK2; *attP* (ØC31): phage attachment site of the ØC31 integrase (= *int* (ØC31)); *xylE*: catechol 2,3-dioxygenase (reporter gene); *tsr*: thiostrepton resistance gene; Thio<sup>R</sup>: Thiostrepton resistance; *bla*: ampicillin resistance gene; *dam*<sup>-</sup>, *dcm*<sup>-</sup>, *hsdS*<sup>-</sup>: methylases-deficient *E. coli* strain; Cml<sup>R</sup>: chloramphenicol resistance; Tet<sup>R</sup>: tetracyclin resistance.

Finally, only one single BGC is similar to the *fev* and *bag* gene clusters in *S. lunaelactis* MM109^T^ as well as in the genomes of 16 other *S. lunaelactis* strains. This makes it extremely unlikely that bagremycins and ferroverdins should be specified by two different BGCs. A plausible pathway for the synthesis of both bagremycins and ferroverdin A from the *fev/bag* cluster is proposed in the Discussion section (see also Figure 8).

### Inactivation and/or duplication of *fevR* affects both ferroverdin and bagremycin production

Of the *S. lunaelactis* strains that were collected from the moonmilk samples, strain *S. lunaelactis* MM91, which does possess a complete *fev/bag* cluster (GenBank: MG708299), failed to produce the ferroverdin-associated green pigmentation when grown in iron-containing media (Figure 6A and 6B).

**Figure 6.**
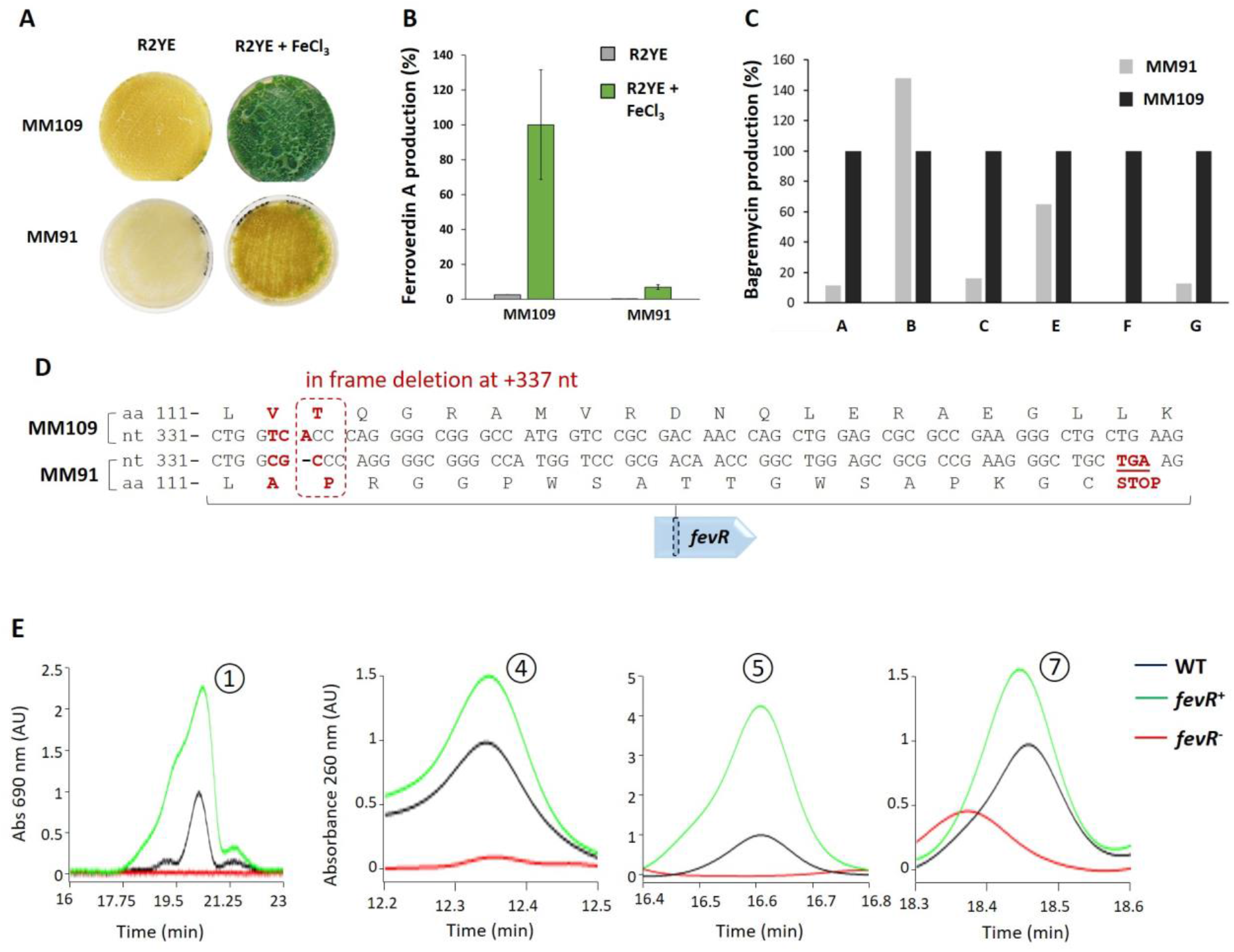
FevR (BagI) is involved in the production of both ferroverdin A and bagremycins. **(A)** Phenotype of *S. lunaelactis* strains MM109^T^ and MM91 grown in media activating (R2YE + FeCl_3_) or repressing (R2YE) the production of ferroverdin A and **(B)** semi-quantitative analysis (HPLC) of ferroverdin A produced by these strains. **(C)** Semi-quantitative analysis (HPLC) of bagremycins produced by *S. lunaelactis* strains MM109^T^ and MM91. **(D)** Identification in the *fev* cluster of *S. lunaelactis* strain MM91 (accession number MG708299) of the in-frame deletion at position +337 nt of *fevR* encoding the SARP-family transcription activator. **(E)** Details of HPLC profiles focused on peaks associated with bagremycin A, bagremycin B, bagremycin F, and ferroverdins from extracts of mutant strains of *S. lunaelactis* MM109^T^ in which *fevR* is inactivated (*fevR*^-^) or with one supplementary copy of *fevR* (*fevR*^+^).

MM91 also showed reduced production of all bagremycins except bagremycin B (Figure 6C). The phenotype of *S. lunaelactis* MM91 allowed a forward genetic approach to assess if the *fev* cluster is indeed responsible for production of both bagremycins and ferroverdins. Genome sequencing and SNP analysis of *S. lunaelactis* strain MM91 revealed a deletion at nucleotide position +337 of SLUN_21380 (*bagI* or *fevR*), which encodes the likely pathway-specific activator of the BGC (Figure 6D). This deletion results in a frameshift leading to a premature stop codon, 17 amino acids downstream of the deletion, and thus in a truncated (and likely inactive) protein.

To ascertain that FevR/BagI is required for both ferroverdin and bagremycin production, the gene SLUN_21380 was first interrupted in strain *S. lunaelactis* MM109^T^ by a thiostreptone resistance cassette. Introduction of plasmid pBDF028 into spores of MM109^T^ allowed to isolate thiostreptone resistant clones in which occurred a single recombination event of a 791 bp internal fragment of SLUN_21380 (*fevR/bagI*). The single cross-over results in a duplication of *fevR*, the first copy lacking the last 86 nt resulting in a truncated FevR protein, and the second copy lacking the promoter region and the first 116 nt. One clone (strain named *fevR*^-^ hereafter) was selected for further analyses. Strain *fevR*^-^ displays loss of the ferroverdin-associated green pigmentation when grown on iron-supplemented R2YE medium. HPLC analysis of the crude extracts of strain *fevR*^-^ confirmed the absence of ferroverdins and bagremycins (Figure 6E). These data suggest that indeed FevR/BagI is the pathway-specific activator for the BGC, and this was further supported by the observation that introduction of an additional copy of *fevR/bagI* into *S. lunaelactis* MM109^T^ resulted in increased production of both ferroverdins and bagremycins (Figure 6E).

### Iron is required for production of *p*-vinylphenyl-3,4-NHBA but not mandatory for bagremycin biosynthesis

If ferroverdins are only detected when iron is supplied in the cultivation media a key question that remains unanswered is whether iron is only required for complexation of the *p*-vinylphenyl-3.4-NHBA monomers (and/or the other monomers required for ferroverdin B and C), or if iron is necessary for the synthesis of *p*-vinylphenyl-3,4-NHBA, the monomer of ferroverdins. An HPLC chromatogram of extracts of *S. lunaelactis* MM109^T^ grown on R2YE and R2YE supplied with 1 mM FeCl_3_ revealed the presence of the *p*-vinylphenyl-3,4-NHBA monomer only when strain MM109^T^ was grown under conditions of high iron (Figure 7A). HPLC fractionation and subsequent UPLC-MS/MS analysis confirmed the presence of a molecular ion species that corresponds to *p*-vinylphenyl-3,4-NHBA (m/z 268.06 [M^-^]) (Figure 7B and 7C). Instead, bagremycins were detected in either conditions (Supplementary Figure S2), suggesting that, in contrast to *p*-vinylphenyl-3,4-NHBA, iron overload is not mandatory for their biosynthesis. Iron is thus not only required for complexation of the three *p*-vinylphenyl-3.4-NHBA ligands with ferrous ion and generate ferroverdin A, but is also necessary for their biosynthesis.

**Figure 7.**
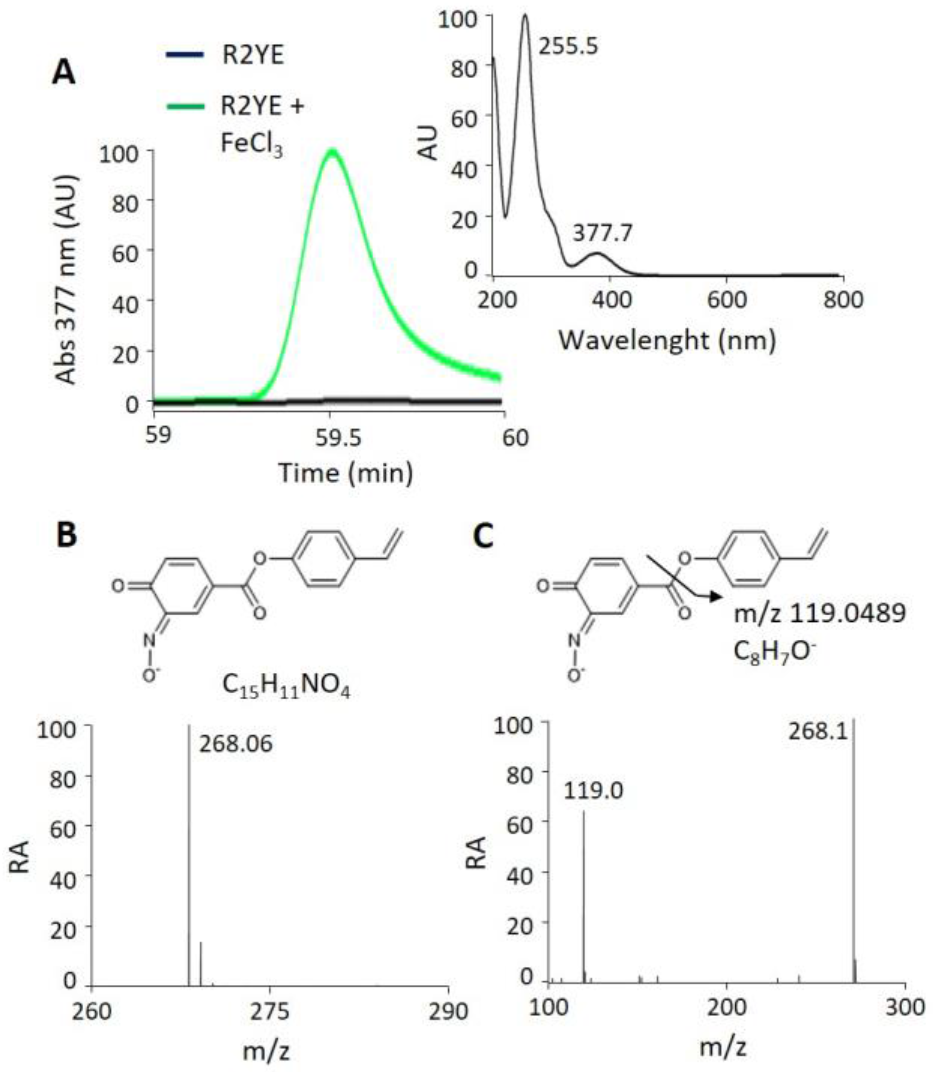
Iron supply is mandatory for production of *p*-vinylphenyl-3-nitroso-4-hydroxybenzoate. **A.** HPLC chromatogram showing the identification of *p*-vinylphenyl-3-nitroso-4-hydroxybenzoate in the extracts of *S. lunaelactis* MM109 grown on R2YE + 1 mM FeCl_3_ (green curve) and the absorbance spectrum from 200 to 800 nm of *p*-vinylphenyl-3-nitroso-4-hydroxybenzoate is shown. Note the absence of the compound when *S. lunaelactis* MM109 is grown on R2YE (black line). **B.** Mass spectrum of the compound present in the chromatographic peak and its proposed structure. **C.** MS/MS spectra of *p*-vinylphenyl-3-nitroso-4-hydroxybenzoate with the proposed fragmentation mechanism.

## Discussion

In this work we provided genomic, genetic and metabolomic evidences that a single BGC (*fev/bag*) is responsible for the synthesis of both ferroverdins and bagremycins. That one single BGC is responsible for the production of several related compounds is not unusual but several features make the case of the *fev/bag* BGC unique. Firstly, the chemical composition of ferroverdins and bagremycins (nitroso-vs amino-aromatic compounds), and secondly, their structural organization (trimers vs monomers) are different while in other BGCs producing multiple compounds these two characteristics are conserved. But what makes this biosynthetic pathway exceptional is that bagremycins and *p*-vinylphenyl-3,4-NHBA, the monomer of ferroverdins, are not produced under the same culture conditions. Indeed, excess of iron supply is mandatory for *p*-vinylphenyl-3,4-NHBA production and for its subsequent complexation to a tri-partite molecule, while the bagremycins are produced under conditions of low iron. Thus, iron is a key molecule that decides between distinct biological activities produced by *S. lunaelactis:* bagremycins display antibacterial, antifungal and anti-proliferative activities presumably in order to inhibit growth of neighboring competing bacteria and fungi and/or to control self-proliferation, while *p*-vinylphenyl-3,4-NHBA production would be only required when iron excess constitutes a main threat in order to limit iron-mediated oxidative damages. The change of the amino-in a nitroso-group in bagremycin allows complexation with iron, which strongly suggests an evolutionary driver for this chemical diversification.

We propose a possible biosynthetic pathway for both bagremycins and ferroverdins (Figure 8) based on the deduced function of the different components of the *bag/fev* clusters, and the current knowledge on the biosynthetic pathway of bagremycins [22, 30, 31, 34]. In addition, as biosynthesis is known for two other types of amino/nitroso-aromatic compounds, namely grixazones (*gri* genes) and 4-hydroxy-3-nitrosobenzamide (*nsp* genes), the information available for these BGCs is useful to predict steps shared by the studied pathways.

**Figure 8.**
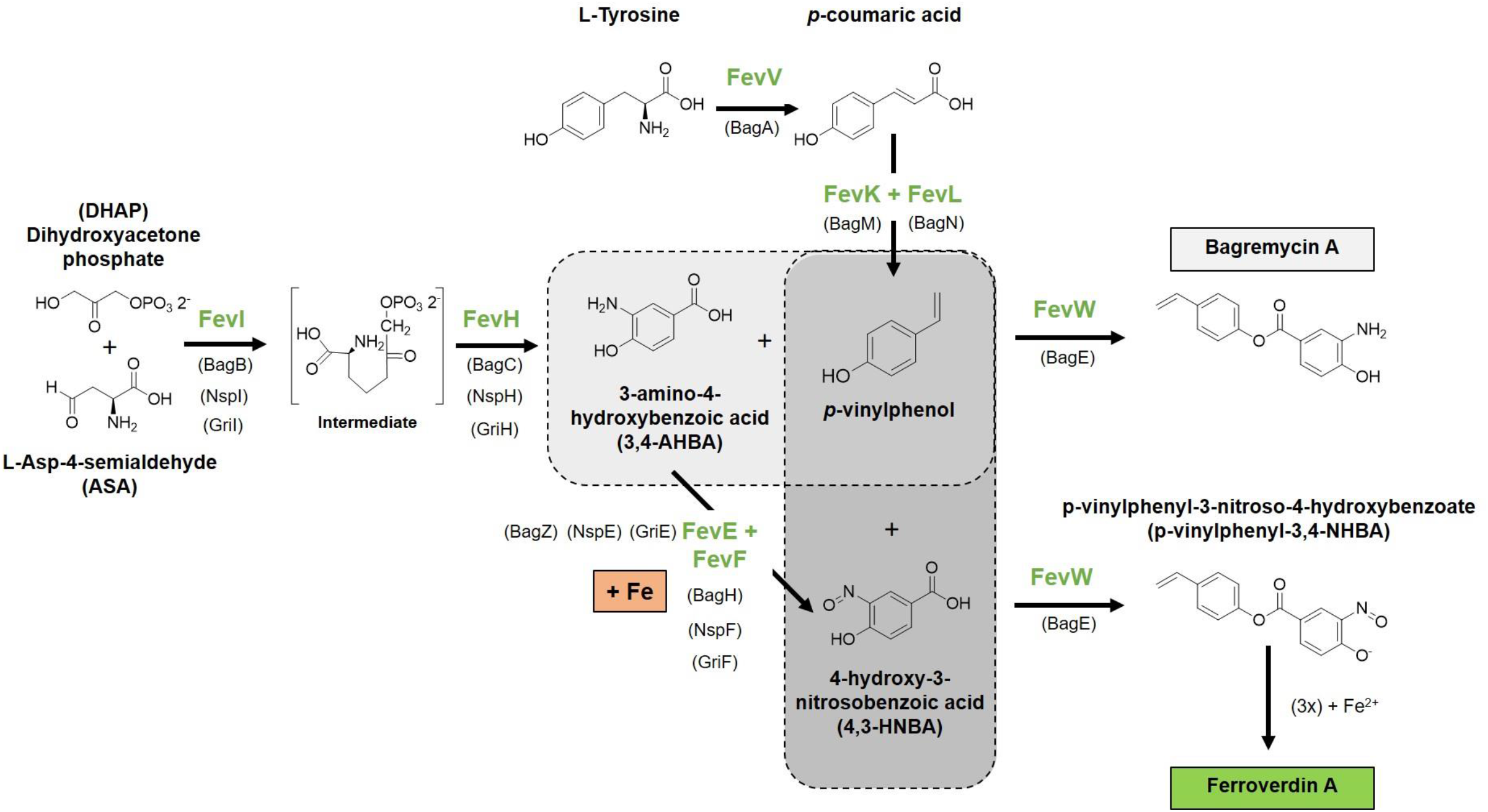
Proposed pathway for bagremycin A and ferroverdin A biosynthesis.

As demonstrated previously, bagremycin A is derived from precursors *p*-vinylphenol and 3-amino-4-hydroxybenzoic acid (3,4-AHBA) [22]. The *p*-vinylphenol is generated in two steps from L-tyrosine which is first deaminated by FevV/BagA in *trans*-coumaric acid, the latter being decarboxylated in *p*-vinylphenol by FevK (BagM) and FevL (BagN). Indeed, FevK and FevL form a UbiX-UbiD system where UbiD is a decarboxylase that requires prenylated FMN as cofactor, which is provided by the flavin prenyltransferase UbiX [35]. Regarding the second precursor 3,4-AHBA, it was demonstrated that it is formed from dihydroxyacetone phosphate (DHAP) and L-aspartic-4-semialdehyde (ASA) through two reactions catalyzed by BagB (FevI) and BagC (FevH) [30]. Condensation of the *p*-vinylphenol and the 3,4-AHBA to form bagremycin A would be mediated by FevW which shows high similarities with proteins of the phenylacetate-CoA ligase family, which catalyze the condensation of two molecules by their carboxillic and alcohol moieties through adenylation.

Regarding the synthesis of *p*-vinylphenyl-3,4-NHBA, the monomer of ferroverdin A and major monomer of other ferroverdins, we propose that it would directly result from precursors *p*-vinylphenol and 4-hydroxy-3-nitrosobenzoic acid (4,3-HNBA) (Figure 7). 4.3-HNBA would be directly generated via the nitrosation of 3,4-AHBA and would be mediated by the copper-containing oxidase FevF (with FevE as chaperone). FevF is a copper-containing oxidase which is orthologous to NspF and GriF, while FevE is a copper chaperone orthologous to NspE and GriE (Table 1). NspE bring two coppers to NspF, which converts 3-amino-4-hydroxybenzamide into 4-hydroxy-3-nitrosobenzamide in *Streptomyces murayamaensis* [32, 36]. Likewise, GriE donates two coppers to GriF, which converts *o*-aminophenol into *o*-quinone imine in *Streptomyces griseus* [37]. Since these FevF/FevE orthologous proteins mediate the oxidation of 3.4-AHBA-like substrates to generate quinone imine moieties which would be consecutively nitrosilated, we suggest that these enzymes are both able to mediate the transformation of 3,4-AHBA to 4,3-HNBA. Condensation of 4,3-HNBA with *p*-vinylphenol by FevW would result in the formation of *p*-vinylphenyl-3-nitroso-4-hydroxybenzoate, the monomer of ferroverdins.

Finally, next to the role of iron as vital element for growth [38, 39], morphogenesis and metabolite production [12, 39, 40], our work reveals an unprecedented complexity of natural product biosynthetic pathways where this element is also an environmental trigger able to change the pattern of natural compounds produced by a single BGC. By challenging the concept that one BGC should produce a single family of molecules, our results also highlight the difficulty to estimate the metabolic potential of an organism based on the genomic information alone.

## Materials and methods

### Strains and culture conditions

All strains used in this study are listed in Table 1. *Escherichia coli* strains were grown in liquid LB or on solid LB medium agar plates and incubated at 37°C. Media used for solid *Streptomyces* cultures were R2YE, SFM, and ISP7 agar plates and incubated at 28°C. When required, antibiotics thiostreptone, kanamycin, nalidixic acid, and ampicillin were added in the culture media. The type strain *S. lunaelactis* MM109^T^ [25] and other *S. lunaelactis* strains were isolated from the cave ‘Grotte des Collemboles’ Collemboles’ (Comblain-au-Pont, Belgium) [25–27]. Media composition and methods associated with *Streptomyces* species were all performed as described in [41].

### Creation of *fevR* knock-out and overexpression mutants in *S. lunaelactis* MM109

In order to assess the effect of *fevR/bagI* on both bagremycin and ferroverdin production we first generated a *S. lunaelactis* MM109 knock-out strain in which *orf* SLUN21380 was interrupted by a single cross-over with a thiostreptone resistance cassette. A 791 bp internal fragment (starting at position +116 nt and lacking the 81 last nt of SLUN21380) was amplified by PCR using primers FevR+118f_XbaI (TC<u>TCTAGA</u>CGAACCAGGTGGTCAGCC) and FevR+909r_*Pst*I (CG<u>CTGCAGG</u>TCGTTCTCCAGGCGC). The PCR product was cloned into the pJET1.2 (Blunt cloning vector, ThermoFisher Scientific) which was named pBDF027 and sequenced for verification of the amplified PCR fragment. This plasmid was then digested by *XbaI* and *Pst*I-HF restriction enzymes (NEB) and the *fevR* insert was subsequently cloned into pSET151 [41] digested with the same enzymes. The resulting plasmid after ligation, named pBDF028 was introduced into *S. lunaelactis* MM109 chromosome by intergeneric conjugative transfer using the methylation-deficient *Escherichia coli* ET12567 containing the plasmid pUZ8002 (ETpUZ) as conjugation partner. After growing *E. coli* ETpUZ to an OD of 0.4 (50 mL culture in LB + chloramphenicol (30 μg/mL) + kanamycin (50 μg/mL) + ampicillin (100 μg/mL)), cells were washed to remove antibiotics and mixed with *S. lunaelactis* spores for mating. Exconjugants selection was carried out on Soy Flour Mannitol (SFM) medium (+ 10 mM MgCl_2_) overlaid with thiostrepton (50 μg/mL) and nalidixic acid (25 μg/mL) after 16 hours of incubation at 28 °C. The single crossover of pBDF028 into the *fevR* locus, was confirmed by PCR and one selected clone was named *fevR*^-^ for further analyses. For complementation, an entire copy of *fevR* (slun21380) including the 217 bp upstream and 360 bp downstream regions was amplified by PCR with primers BDF53 (GATCTAGAAAGCTTGGCTCTGTCCAGTGAGACATCC) and BDF54 (GCGAATTCGTACTCGATGTCACCCGCC) to obtain a 1570 bp fragment flanked by EcoRI and XbaI restriction sites. The PCR product was first cloned into a pJET1.2 which was named pBDF019. Using EcoRI-HF and XbaI enzymes, the 1570 bp fevR-containing fragment was cloned into a pSET152 [41] integrative vector leading to pBDF021. Conjugation was performed with spores (10^8^) of *S. lunaelactis* MM109 containing pBDF028 on SFM (+ 10 mM MgCl_2_) thiostrepton (50 μg/mL) overlaid with apramycin and nalidixic acid. Integration of the plasmid was confirmed by PCR with pSET_forward (GAGCGGATAACAATTTCACACAGGA) and BDF53 or pSET_reverse (CGCCAGGGTTTTCCCAGTCACGAC) and BDF54 primers.

### Compound extraction and analysis by High Pressure Liquid Chromatography (HPLC)

Compound extraction and HPLC analysis were mainly performed as described previously [26]. All *S. lunaelactis* strains were cultured for 10 days on different solid media on Petri dishes (90 mm, 25 ml of medium) and incubated at 28 C°. The solid cultures were cut into small pieces (about 0.5 × 0.5 cm) and then mixed overnight with an equal volume (25 ml per plate) of ethyl acetate. The mixture was centrifuged (20 min at 4000 rpm) and the supernatant was evaporated (25°C at 210 rpm) on a rotary evaporator (IKA RV10 digital, VWR, Radnor, PA, USA). The dried crude extract was resuspended in 1 ml of acetonitrile for further analyses. The full extract was then fractionated by HPLC (Waters, Milford, MA, USA) using a Waters 2695 Separations Module (Alliance) with a Waters 2998 Photodiode Array Detector coupled to a Waters Fraction Collector WFC III. The extracts were analyzed on a Luna Omega PS C18 column (2.1 mm × 150 mm, 5 μm particle size, 100 Å, Phenomenex) at a column temperature of 40 °C. Extract separation was achieved by increasing the acetonitrile (Barker, HPLC far UV grade)/water (milliQ filtrated on 0.22 μm) + 0.05% trifluoroacetic acid (TFA, Sequencing grade; Thermo Fisher Scientific, San Jose, CA, USA), ratio (from 0 to 100% of acetonitrile during 80 min) at a 450 μl/min flow rate. Online UV absorption measurement was performed from 190 to 800 nm. Data were analyzed using software Empower 3 (Waters, Milford, MA, USA), and Xcalibur v2.2 (Thermo Fisher Scientific, San Jose, CA, USA). Fractions were subsequently tested for antibacterial activities against *Staphylococcus aureus* (ATCC25923) by disk diffusion assay as describe previously [26].

### Compound identification by Ultra-Performance Liquid Chromatography-Tandem Mass Spectrometry (UPLC-MS/MS)

UV-VIS spectra absorbance were obtained by analytical HPLC RP-C18 analyses. HRESIMS data were acquired on a Q Exactive Plus hybrid Quadrupole-Orbitrap Mass Spectrometer (Thermo Fisher Scientific, San Jose, CA, USA). Briefly, compounds were separated by reverse-phase chromatography using Ultra Performance Liquid chromatography (UPLC IClass, Waters) using a Nucleodur C18ec column (2.0 mm × 150 mm, 5 μm particle size, Macherey-Nagel). Elution was achieved by increasing the acetonitrile/water (milliQ filtrated on 0.22 μm) + 0.05% trifluoroacetic acid (for positive ionization mode) ratio (from 0 to 62.5 % during 30 min, then from 62.5 to 100 % during 8 min) at a 300 μl/min flow rate. For the negative ionization mode elution was achieved by increasing the acetonitrile/water (milliQ filtrated on 0.22 μm) + 0.1% formic acid ratio (from 0 to 62.5 % during 30 min, then from 62.5 to 100% during 8 min) at a 300 μl/min flow rate on a Luna Omega PS C18 column (2.1 mm × 150 mm, 5 μm particle size, 100 Å pore size, Phenomenex). On-line UV absorption measurement was performed at 210 and 265 nm and the chromatography system was finally coupled to a Q Exactive Plus hybrid Quadrupole-Orbitrap Mass Spectrometer (Thermo Fisher Scientific, San Jose, CA, USA), operated in positive ion mode (for bagremycins) and in negative mode (for ferroverdins and bagremycins), and programmed for data-dependent acquisitions. Survey scans were acquired at mass resolving power of 140,000 FWHM (full width at half maximum) from 100 to 1500 m/z (1 × 106 ions accumulation target). The five most intense ions were then selected to perform MS/MS experiments by Higher Energy Collision Dissociation (HCD) fragmentations using stepped normalized collision energy (NCE; 21,2; 25; 28) within 2 amu isolation windows (resolution 17500, 1 × 105 ions accumulation target). A dynamic exclusion was enabled for 10 s. Data were analyzed using Xcalibur v2.2 (Thermo Fisher Scientific, San Jose, CA, USA).

Each compounds was identified by its exact mass, isotope pattern, observation of the MS/MS spectra obtained on the molecular ion fragmentation, and the UV-VIS absorbance spectra. For each molecule a fragmentation pathway was proposed (see Supplementary Figure S3).

### Bioinformatics

The complete genome sequence of *S. lunaelactis* strains was obtained by Illumina HiSeq, and MiSeq (Illumina, CA, USA) technologies as described previously [28]. The two sequencing techniques were performed at the GIGA-Research Center (Liège University, Belgium), and at the Luxembourg Institute of Science and Technology (Belvaux, Luxembourg), respectively. Genomes were assembled as described previously and using the complete genome sequence of the type strain *S. lunaelactis* MM109^T^ as template [29].

Phylogeny analyses were performed with sequences of homolog proteins from the *fev (S. lunaelactis* MM109, S. sp. WK-5344), *bag* (S. sp. Tü 4128), *gri (S. griseus*), and *nsp* (*S. murayamaensis*) clusters. Proteins were aligned with MAFFT (v7.273, localpair mode) and alignments were trimmed and internal gaps removed. Phylogenetic inference was deduced with the maximum likelihood method as implemented in RAxML (v8.1.17, rapid bootstrapping mode with 1000 replicates, GTR+I+G evolutionary model).

## Acknowledgements

LM and MM work was supported by a Research Foundation for Industry and Agriculture (FRIA) grant. AN work is supported by a First Spin-off grant from the Walloon Region (Grant number: 1510530; FSO AntiPred). SR is a Maître de Recherche at Belgian Fund for Scientific Research (F.R.S-FNRS). GVW acknowledges NACTAR grant 16439 by the Netherlands Organization for Scientific Research (NWO)

## Supplementary data

**Supplementary Figure S1.**
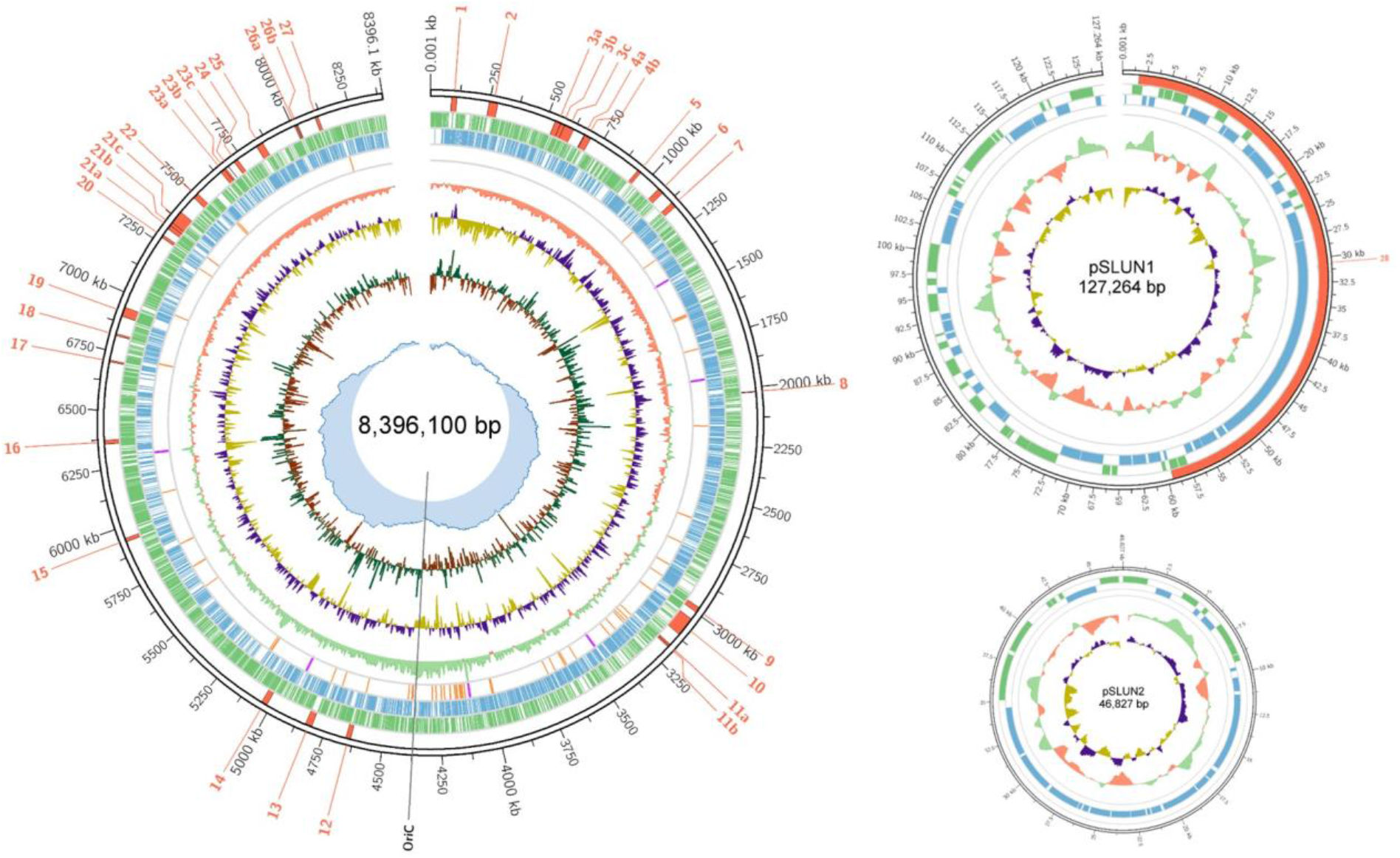
Schematic representation of *S. lunaelactis* MM109^T^ chromosome and its two extrachromosomal plasmids. **A**. Chromosome. From outside to insides, the concentric tracks represent: nucleotide position; predicted BGCs with associated labels; CDSs on the forward strand; CDSs on the reverse strand; tRNA (orange) and rRNA (magenta) genes; per-base read coverage (green and red for above and below average, window size = 10,000; base step size = 1000); GC content (GC %, window size = 10,000; base step size = 1000); GC skew ([G+C]/[G-C], window size = 8,396; base step size = 839); cumulative GC skew. The position of the predicted origin of replication (OriC) is indicated. **B**. Linear plasmid pSLUN1. C. Circular plasmid pSLUN2. The same track coloring applies for the plasmids. (The coverage and GC content sliding averages have a window size of 1,000 bp and a base step size of 100 bp).

**Supplementary Figure S2.**
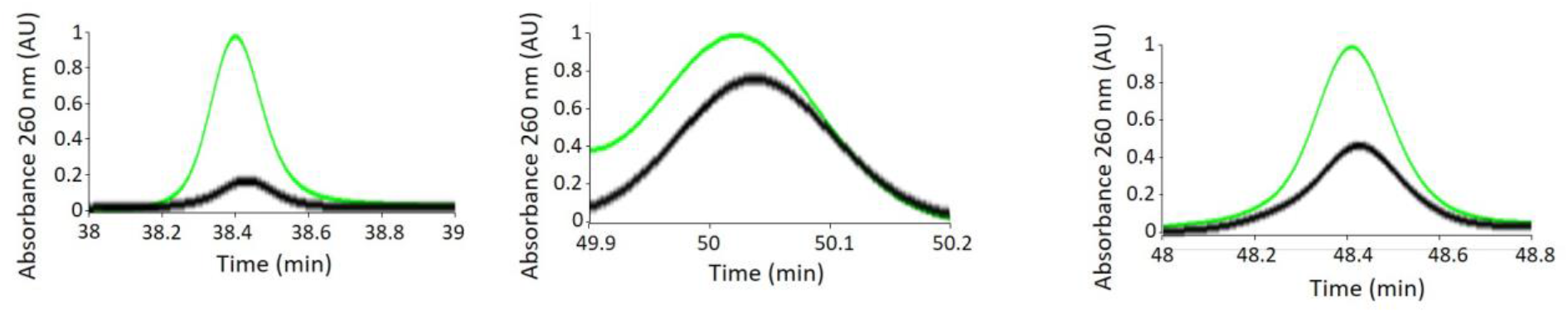
Identification of bagremycin A (left graph), bagremycin B (middle graph), and bagremycin G (right graph) in the culture extracts of *Streptomyces lunaelactis* MM109 of grown on R2YE (black curve) and R2YE + 1 mM FeCl_3_ (green curve).

**Supplementary Figure S3.**
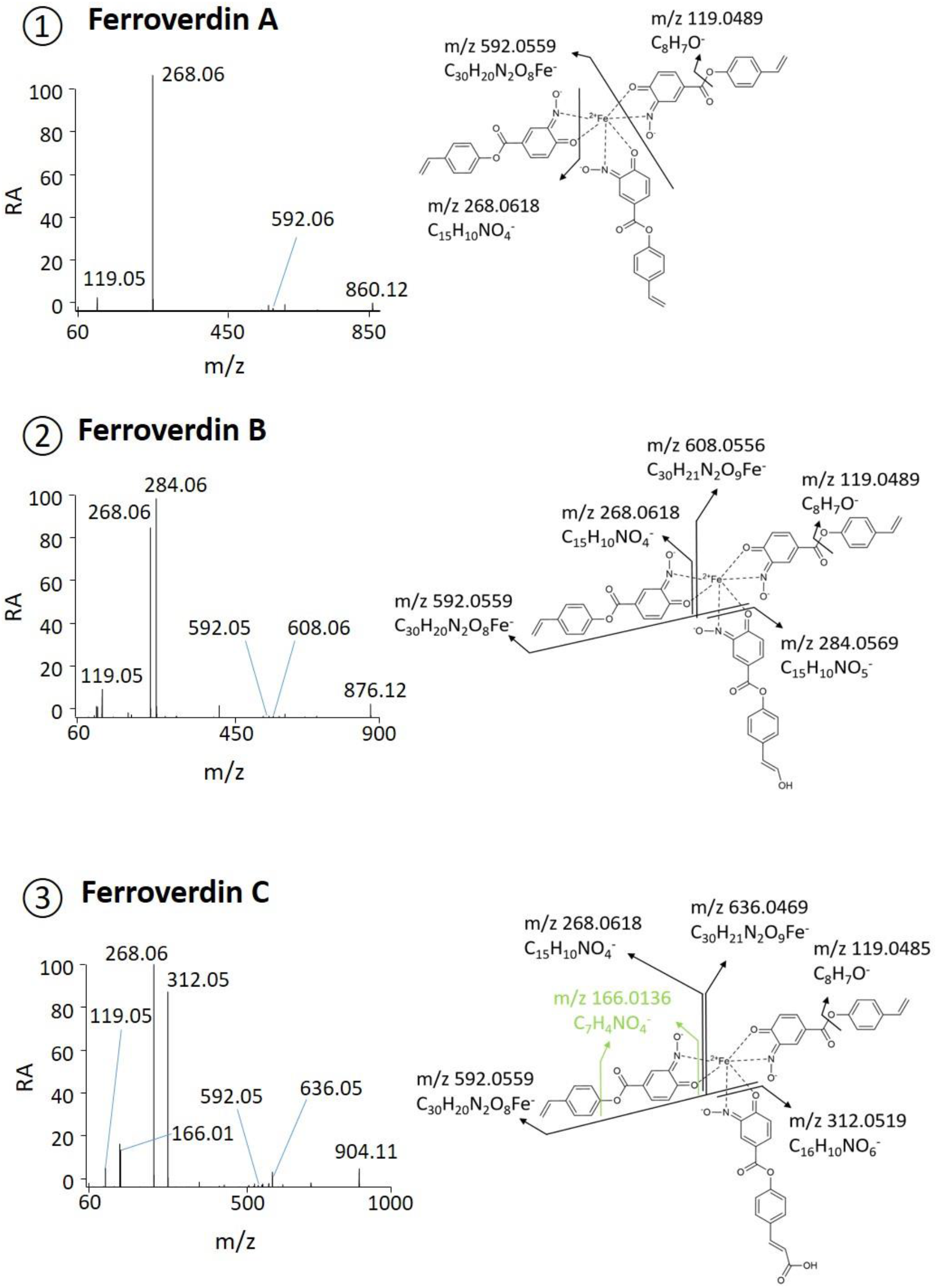
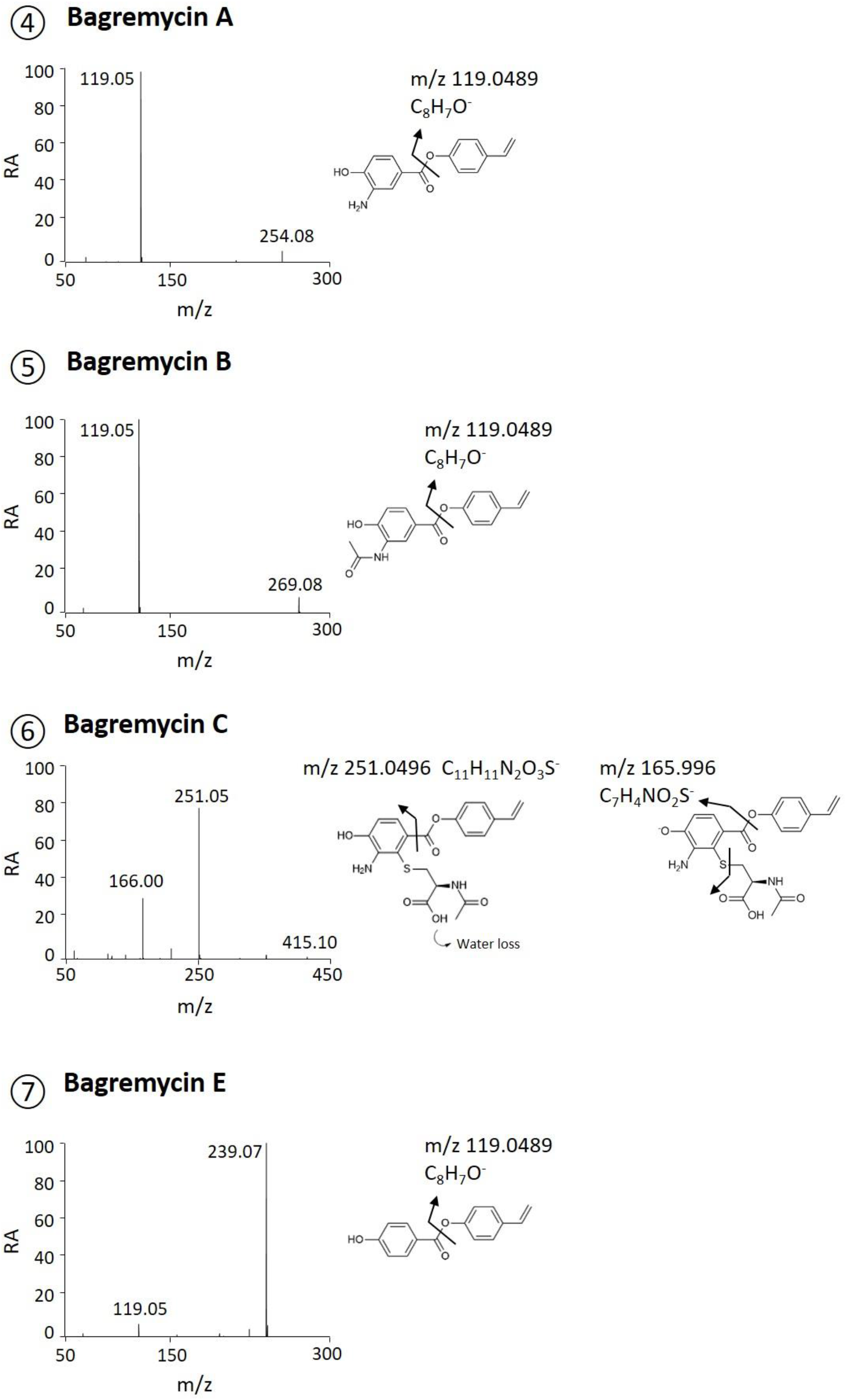
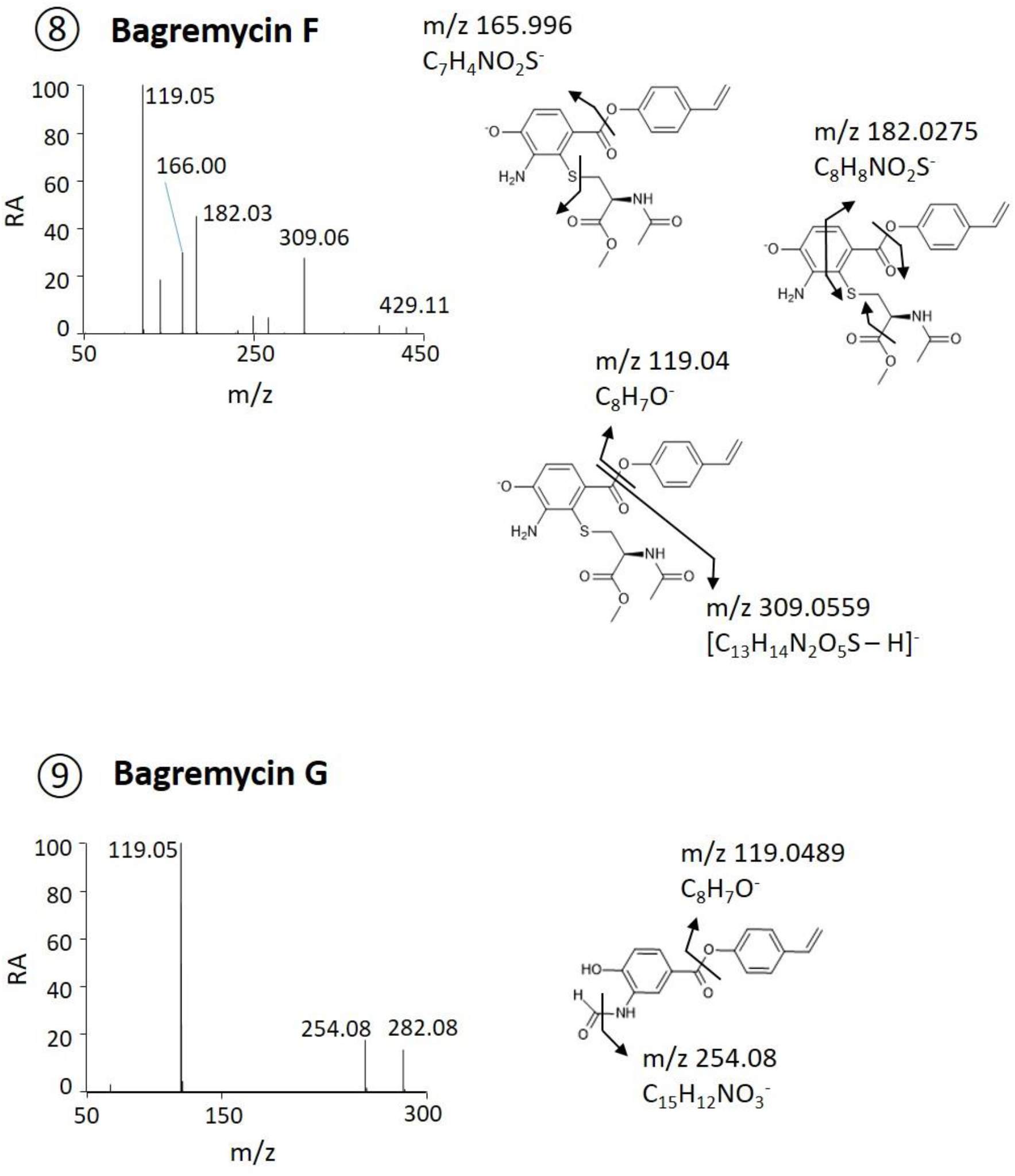
ESI(neg)–MS/MS spectra and proposed fragmentation mechanisms for each deprotonated molecule identified in this study. Fragmentation mechanisms of ferroverdins (compounds 1 to 3) obtained by HCD fragmentation of the molecular ions (M-) 860.12, 876.11 and 904.11 respectively. Fragmentation mechanisms of bagremycins (compounds 4 to 9) were obtained by HCD fragmentation of the molecular ions (M-H)^-^ 254.08, 269.09, 415.10, 239.07, 429.11, and 282.08, respectively.

